# Circulating selenium and prostate cancer risk: a Mendelian randomization analysis

**DOI:** 10.1101/223248

**Authors:** James Yarmolinsky, Carolina Bonilla, Philip C Haycock, Ryan JQ Langdon, Luca A Lotta, Claudia Langenberg, Caroline L Relton, Sarah J Lewis, David M Evans, the PRACTICAL consortium, George Davey Smith, Richard M Martin

## Abstract

In the Selenium and Vitamin E Cancer Prevention Trial (SELECT), selenium supplementation (causing a median 114 μg/L increase in circulating selenium) did not lower overall prostate cancer risk, but increased risk of high-grade prostate cancer and type 2 diabetes. Mendelian randomization analysis uses genetic variants to proxy modifiable risk factors and can strengthen causal inference in observational studies. We constructed a genetic risk score comprising eleven single-nucleotide polymorphisms robustly (*P*<5x10^−8^) associated with circulating selenium in genome-wide association studies. In a Mendelian randomization analysis of 72,729 men in the PRACTICAL Consortium (44,825 cases, 27,904 controls), 114 μg/L higher genetically-elevated circulating selenium was not associated with prostate cancer (OR: 1.01; 95% CI: 0.89-1.13). Concordant with findings from SELECT, selenium was weakly associated with advanced (including high-grade) prostate cancer (OR: 1.21; 95% CI: 0.98-1.49) and type 2 diabetes (OR: 1.18; 95% CI: 0.97-1.43; in a type 2 diabetes GWAS meta-analysis with up to 49,266 cases, 249,906 controls). Mendelian randomization mirrored the outcome of selenium supplementation in SELECT and may offer an approach for the prioritization of interventions for follow-up in large-scale randomized controlled trials.

---

The development of interventions to prevent cancer requires robust causal knowledge, but few observational epidemiological claims are replicated in randomized controlled trials (RCTs) and some trial results are in the opposite direction to those seen observationally (i.e. causing harm) (1, 2). Such failures to translate observational data into effective cancer prevention interventions arise in part because the inherent limitations of observational research - confounding, reverse causation, and measurement error - preclude confident causal inference.

Mendelian randomization uses genetic variants as instruments (i.e. proxies) to assess whether a potential intervention target (e.g. risk factor, molecular intermediate, or gene product) has a causal effect on a disease outcome in a non-experimental (observational) setting (3). The principle of Mendelian randomization is that analysis of groups defined by common genetic variants is analogous to that of intention-to-treat analysis in an RCT. This is based on the independent assortment of genetic variants at meiosis which should allow for genotype at a population level to be largely independent of later environment and lifestyle factors (Figure 1). Using genetic variants as instruments to proxy intervention targets means that analyses should be less susceptible to confounding by environmental factors that typically distort observational associations. Furthermore, genotypes are typically measured with little error, represent life-long exposure, and are not subject to reverse causation because disease status cannot influence one’s germline genotype.

**Figure 1.**
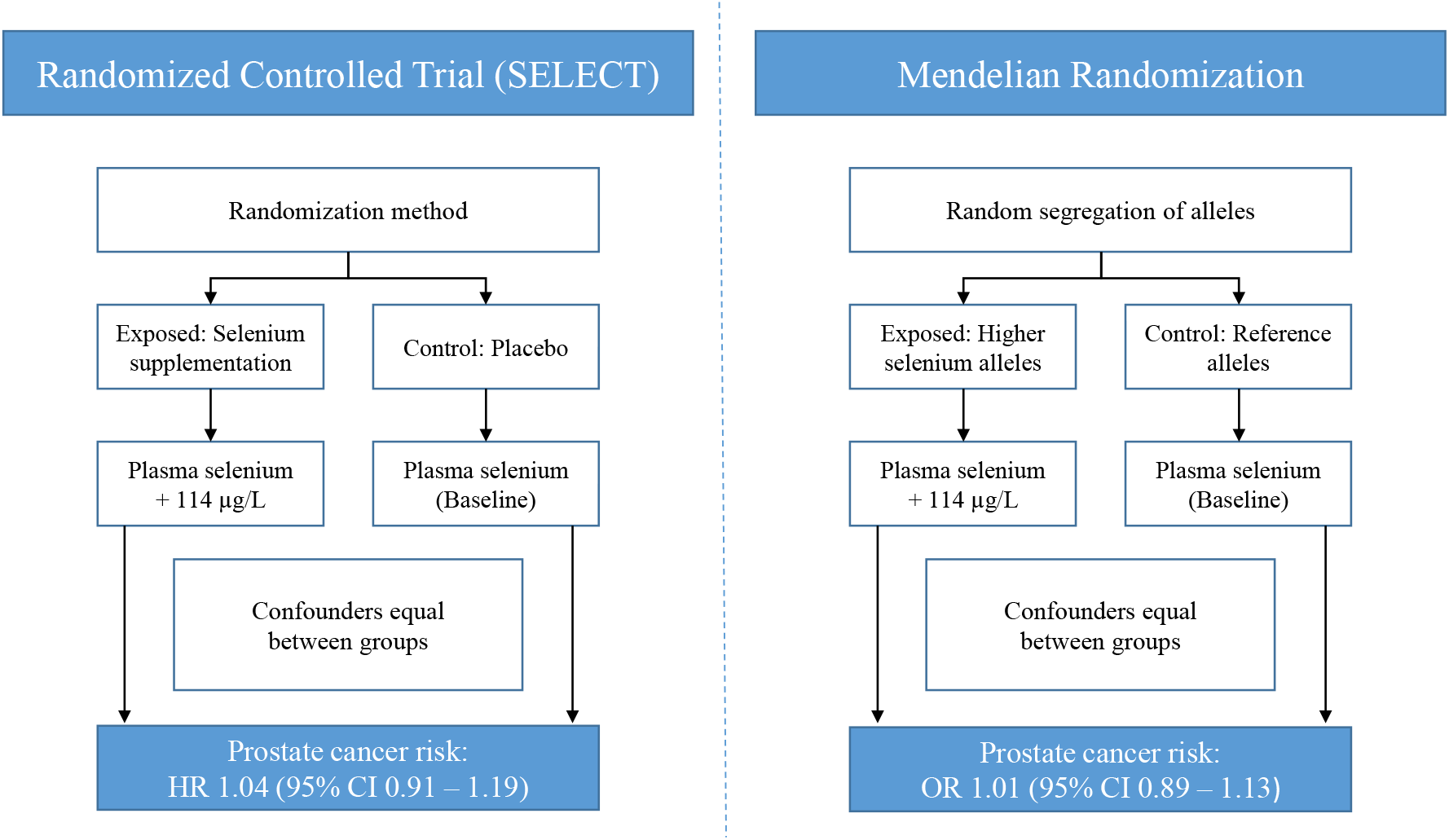
Schematic comparison of a Randomized Controlled Trial (SELECT) to a Mendelian randomization analysis. In an RCT, individuals are randomly allocated to an intervention or control group (In SELECT, 200 μg/d selenium [114μg/L increase in blood selenium] or placebo). If the trial is adequately sized, randomization should ensure that intervention and control groups are comparable in all respects (e.g. approximately equal distribution of potential confounding factors) except for the intervention being tested. In an intention-to-treat analysis, any observed differences in outcomes between intervention and control groups can then be attributed to the trial arm to which they were allocated. In a Mendelian randomization (MR) analysis, alleles that influence levels of a trait of interest are randomly allocated at conception (In MR, the additive effects of selenium-raising alleles at eleven SNPs were scaled to mirror a 114μg/L increase in blood selenium). Groups defined by genotype should be comparable in all respects (e.g. distribution of both genetic and environmental confounding factors) except for their exposure to a trait of interest. Any observed differences in outcomes between groups defined by genotype can then be attributed to differences in life-long exposure to the trait of interest under study. Mendelian randomization is an application of the technique of instrumental variable (IV) analysis. In order for a genetic variant (or a multi-allelic genetic risk score) to be used as an IV, three key assumptions must be met: 1) The instrument must be reliably associated with the exposure of interest, 2) the instrument should be independent of other factors affecting the outcome (confounders), 3) the instrument should only affect the outcome through the exposure of interest (known as the exclusion restriction criterion).

An advantage of Mendelian randomization is that implementation does not require access to individual level data or trait measurements in all samples: it can be implemented using information on genetic variant-exposure and genetic variant-outcome associations obtained from separate samples, which greatly increases the scope of the approach (“two-sample Mendelian randomization”) (4, 5). Two-sample approaches using summary data can be used in an efficient and cost-effective manner to screen hundreds of potential intervention targets for causal relationships with cancer, without having to expensively measure these targets within individual cancer collections, and is made increasingly possible by the rapid increase in genome-wide association studies (GWAS) over the last decade; there are currently over 2000 published GWAS for over 1500 unique traits (https://www.genome.gov/gwastudies/).

The largest ever prostate cancer prevention trial (SELECT, N=35,533) was designed to examine whether daily supplementation with selenium, vitamin E, or both agents combined, could prevent prostate cancer (6). It was abandoned at a cost of $114 million because of lack of efficacy compounded by possible carcinogenic (increased rates of high-grade [Gleason score ≥ 7] prostate cancer) and adverse metabolic effects (increased rates of diabetes) of the interventions (6, 7). We investigated whether Mendelian randomization could have predicted the results of the SELECT trial observed for selenium in a two-sample Mendelian randomization study of 72,729 individuals of European descent from the PRACTICAL (Prostate Cancer Association Group to Investigate Cancer Associated Alterations in the Genome) consortium (8).

We obtained summary GWAS statistics from analyses on 44,825 prostate cancer cases and 27,904 controls of European descent from 108 studies in PRACTICAL. Summary statistics were also obtained from analyses on 6,263 advanced prostate cancer cases (defined as Gleason ≥8, prostate-specific antigen >100 ng/mL, metastatic disease (M1), or death from prostate cancer) and 27,235 controls (summary data on high-grade prostate cancer alone was not available). All studies in PRACTICAL have the relevant Institutional Review Board approval from each country, in accordance with the Declaration of Helsinki. Genotyping of PRACTICAL samples was performed using an Illumina Custom Infinium genotyping array (OncoArray), designed for the OncoArray consortium and consisting of ~570,000 single nucleotide polymorphisms (SNPs) (9). All SNPs with a poor imputation quality (r^2^<0.30), a minor allele frequency of <1%, a call rate of <98%, or evidence of violation of Hardy-Weinberg equilibrium (*P*<10^−7^ in controls or *P*<10^−12^ in cases) were removed.

To analyze the effect of selenium on type 2 diabetes we used summary GWAS data from analyses in up to 49,266 type 2 diabetes cases and 249,906 controls of European descent obtained from a meta-analysis of the DIAbetes Genetics Replication And Meta-analysis (10), EPIC-InterAct (11), and UK Biobank studies (12). Methods for this meta-analysis have been published previously (13).

A genetic risk score to proxy for circulating selenium levels was constructed by obtaining SNPs shown to robustly (*P*<5x10^−8^) associate with selenium concentrations in a meta-analysis of blood and toenail selenium GWAS (14, 15). Of twelve selenium SNPs identified, one (rs558133) was not available in PRACTICAL and thus eleven SNPs were used as genetic instruments for overall and advanced prostate cancer analyses. As sensitivity analyses, we also constructed a restricted genetic risk score using only SNPs robustly (*P*<5x10^−8^) associated with selenium in a GWAS of blood selenium that were replicated (*P*<0.05) in subsequent independent studies (14–16). For these analyses, of five selenium SNPs initially identified, one (rs6859667) was not available in PRACTICAL and thus four SNPs were used as instruments for both prostate cancer analyses. All SNPs utilised for primary and sensitivity analyses and their corresponding ENSEMBL-mapped gene(s) (17) are presented in the footnote to Table 1.

**Table 1.** Comparison of the effect of 114 ug/L selenium on overall prostate cancer, high-grade / advanced prostate cancer, and type 2 diabetes in SELECT and Mendelian randomization

|  | SELECT<br>HR (95% CI) | Mendelian randomization<br>OR (95% CI) |
| --- | --- | --- |
| Overall prostate cancer | 1.04 (0.91 to 1.19) | 1.01 (0.89 to 1.13) |
| High-grade/advanced prostate cancer* | 1.21 (0.97 to 1.52) | 1.21 (0.98 to 1.49) |
| Type 2 diabetes | 1.07 (0.87 to 1.18) | 1.18 (0.97 to 1.43) |

HR= Hazard Ratio; OR= Odds Ratio; CI= Confidence Interval.
\*“High-grade prostate cancer” pertains to SELECT results and “Advanced prostate cancer” pertains to Mendelian randomization results.

We constructed a correlation matrix to estimate correlation between SNPs with reference to the HapMap 3 (release 2) dataset. The causal effect of our genetic risk score on overall prostate cancer, advanced prostate cancer, and type 2 diabetes was then examined using a maximum likelihood approach that takes into account moderate correlations between genetic variants (18). To compare the causal odds ratios from Mendelian randomization with the hazard ratios from SELECT, we estimated the causal odds ratios per 114μg/L genetically increased circulating selenium, to match the measured pre- versus post-intervention blood selenium differences between supplementation and control arms in SELECT (6). All statistical analyses were performed using R version 3.0.2.

In Mendelian randomization analyses, a 114μg/L increase in genetically-elevated blood selenium was not associated with overall prostate cancer risk (OR: 1.01, 95% CI: 0.89–1.13, *P*=0.93) (Table 1). Genetically elevated selenium was weakly associated with advanced prostate cancer (OR: 1.21, 95% CI: 0.98-1.49, *P*=0.07) and type 2 diabetes (OR: 1.18, 95% CI: 0.97-1.43, *P*=0.11). Results for overall prostate cancer, advanced prostate cancer and type 2 diabetes were robust to sensitivity analyses employing a restricted genetic risk score (ORs were 0.95 [95% CI 0.80-1.14], 1.09 [0.81-1.47] and 1.23 [0.99-1.53]), respectively).

Limitations of our analysis include that we were only able to directly examine the effect of genetically elevated selenium on advanced prostate cancer and not high-grade prostate cancer per se (summary estimates from PRACTICAL were available only for a composite “advanced” disease classification), thus preventing direct comparison to results with SELECT. Additionally, our selenium SNPs were also associated with betaine, a putative risk factor for type 2 diabetes (19), which could introduce horizontal pleiotropy (genetic variants influencing an outcome through a different biological pathway from the exposure under investigation) into our analyses and thus invalidate instrumental variable assumptions (see footnote of Figure 1). Though we suspect that the association of these SNPs with both selenium and betaine reflects the effect of selenium on betaine in the methionine cycle (20, 21) and, consequently, that both selenium and betaine likely influence type 2 diabetes risk through the same biological pathway, we cannot rule out the possibility that at least part of a putative effect of selenium SNPs on type 2 diabetes risk is through an alternate biological pathway involving betaine.

We conclude that in contrast to findings from some (22–27) but not all prospective epidemiological studies (28, 29), our Mendelian randomization analysis using publicly available GWAS data did not find strong evidence for a causal effect of selenium on overall prostate cancer. Consistent with SELECT, we found weak evidence of a positive effect of genetically elevated selenium on advanced prostate cancer. In agreement with SELECT, we also found weak evidence of a positive effect of genetically elevated selenium on type 2 diabetes risk. The alignment of Mendelian randomization with SELECT estimates mirrors the concordance of Mendelian randomization findings with other large, phase III trials; including adverse effects of elevated LDL cholesterol on coronary heart disease (30, 31) and statin use with type 2 diabetes (32, 33); and null effects of secretory phospholipase A(2)-IIA with cardiovascular disease (34–36) and HDL cholesterol on risk of myocardial infarction (37–40). Mendelian randomization may serve as an important time-efficient and inexpensive first step in predicting both the efficacy and possible adverse effects of an intervention prior to the design of a randomized controlled trial.

## Abbreviations

Randomized controlled trial (RCT), genome-wide association study (GWAS), Selenium and Vitamin E Cancer Prevention Trial (SELECT), Prostate Cancer Association Group to Investigate Cancer Associated Alterations in the Genome (PRACTICAL), single nucleotide polymorphisms (SNPs)

## Acknowledgments

We are grateful to the contribution of data from EPIC-InterAct Investigators. The InterAct Study is led by Professor Nick Wareham at the MRC Epidemiology Unit, Cambridge, UK; InterAct funding was provided by the EU FP6 programme (grant number LSHM_CT_2006_03 7197).

